# Novel zebrafish mutants reveal new roles for Apolipoprotein B during embryonic development and pathological conditions

**DOI:** 10.1101/2021.04.01.437990

**Authors:** Hanoch Templehof, Noga Moshe, Inbal Avraham-Davidi, Karina Yaniv

**Author notes:** Corresponding Author, Karina Yaniv, Department of Biological Regulation, Weizmann Institute of Science, Rehovot, 76100, Israel.

## Abstract

Apolipoprotein B (ApoB) is the primary protein of chylomicrons, VLDLs and LDLs and is essential for their assembly. Defects in ApoB synthesis and secretion result in several human diseases, including abetalipoproteinemia and familial hypobetalipoproteinemia. Conversely, high levels of APOB in plasma are associated with increased risk for coronary heart disease and atherosclerosis.

The involvement of APOB in lipid metabolism and atherogenesis prompted the generation of several mutant mice. However, as APOB is required for supplying nutrients to the developing embryo, *ApoB* null mice are embryonic lethal, thereby precluding the study of the roles of this protein during development.

Here, we established novel zebrafish mutants for two *apoB* genes: *apoBa* and *apoBb.1*. Double-mutant embryos display clear hallmarks of human hypolipidemia-related diseases, including intestinal defects and fatty liver, as well as profound vascular defects. We further use these models to identify the domains within ApoB responsible for its functions. By assessing the ability of different truncated forms of human APOB to rescue the mutant phenotypes, we demonstrate the benefits of this model for prospective therapeutic screens. Overall, our novel zebrafish models uncover new functions of ApoB in organ development and morphogenesis and shed new light on the mechanisms underlying hypolipidemia-related diseases.

## Introduction

Cardiovascular diseases (CVDs) take a huge toll on the world population. An estimated 18 million people die each year from CVDs, representing 31% of all global deaths. Furthermore, CVDs are projected to remain the single leading cause of death in the Western world. Elevated levels of LDL are widely recognized as a cardiovascular risk factor, and abundant data point to chemically modified LDL and its apoprotein- apolipoprotein B (ApoB) as triggers of most of the features of the pathobiology of atherosclerosis.

ApoB is the primary structural protein of chylomicrons, VLDLs and LDLs, and is essential for their assembly^1^. Two APOB isoforms- APOB100 and APOB48- are present in humans and mice, which are encoded by a single gene and generated by RNA editing^2^. Defects in ApoB synthesis and secretion result in several human diseases, including abetalipoproteinemia (ABL) and familial hypobetalipoproteinemia (FHBL1)^3,4^, a human autosomal dominant disorder caused by a missense mutation in the *APOB100* gene. In contrast, high levels of APOB in plasma are associated with an increased risk for coronary heart disease and atherosclerosis^1,5^.

In addition to the involvement of ApoB in CVDs, ApoB-related dyslipidemia is also linked to another global epidemic- non-alcoholic fatty liver disease (NAFLD), which affects around 20%-30% of the adult population in Western countries^6^. NAFLD results in steatosis, the accumulation of lipid in hepatocytes^7^, which can lead to liver malfunction, cirrhosis and, in some cases, to hepatocellular carcinoma^8^. Defects in production and/or secretion of ApoB lipoproteins are highly linked to hepatic steatosis^9^. In particular, the impaired synthesis of VLDL observed in FHBL patients, which results in triglyceride accumulation in the liver, represents one among many causes of NAFLD in humans^10^. In addition to the involvement of APOB in atherogenesis and its important role in lipid metabolism, a large body of data accumulated during the past years, has revealed new roles for lipoproteins as signaling mediators in various cell types, operating at different levels and through various classic and non-classic mechanisms^11^. This, has prompted the generation of several genetically modified mouse models^12,13^. However, as APOB is expressed in the yolk sac during early embryonic development, where it facilitates the supply of nutrients to the developing embryo^14^, ApoB knockout mice die at gestational stage 9.5-10.5, thereby precluding the study of the roles of this protein during embryogenesis.

The close relationship between zebrafish and human apolipoprotein expression and function^15,16^ makes the former an attractive animal model for studying the repertoire of ApoB functions during embryonic development. In addition, the well-documented ability of zebrafish to develop hyperlipidemia^17,18^ and lipoprotein oxidation^19,20^ motivated the study of the cellular and molecular mechanisms linking lipoproteins and CVD in this animal. Here, we established novel mutants for two zebrafish *apoB* genes: *apoBa* and *apoBb.1*. We find that double mutant embryos display lipid accumulation in hepatocytes, a common hallmark of FHBL1 and NAFLD. In addition, we detect early developmental defects, including abnormal liver laterality, decreased numbers of goblet cells in the gut and impaired angiogenesis in double *apoB* mutants. Interestingly, we find altered Notch signaling to underlie some of the observed phenotypes. To identify the domains within ApoB associated with the anti-angiogenic function, we assessed the ability of truncated forms of the human APOB-APOB25 and APOB34-, to rescue the zebrafish mutant phenotypes. Interestingly, only mutant embryos injected with APOB34 displayed a significant rescue of the vascular phenotypes suggesting that the anti-angiogenic activity of ApoB is contained between the 25 to 34-N terminus remnant of the ApoB protein. Altogether, these results uncover previously unappreciated functions of ApoB during early organ development and highlight the potential of the novel *apoB* zebrafish mutants as models for studying human pathologies associated with hypolipidemia, as well as for related drug screens.

## Results

### Generation of zebrafish *apoB* mutants

Three *apoB* genes are detected in the zebrafish genome: *apoBa*, *apoBb.1* and *apoB.2*^16,21^. 2 days post-fertilization (dpf) embryos displayed strong expression of *apoBa* and *apoBb.1* mRNA in the yolk syncytial layer (YSL), whereas *apoBb.2* expression was undetectable (Figure 1A,B,E). At 6 dpf, *apoBa* expression became restricted to the liver (Figure 1C, arrowhead), and *apoB.1* was detected mostly in the YSL and the intestine (Figure 1D). To investigate the role of ApoB during embryonic development, we generated zebrafish carrying mutations in the *apoBa* and *apoBb.1* genes. sgRNAs targeting exon 5 of the *apoBb.1* gene (Supplemental Figure 1A) were injected along with *cas9* mRNA into one-cell stage *Tg(fli1:eGFP)*^22^ embryos. F0 injected fish were raised to adulthood and screened for germline transmission. We identified two different mutations, including an 18-bp insertion containing a premature in-frame STOP codon (Supplemental Figure 1A), which was used throughout this study. To target the *apoBa* gene, we used the TALEN technology. mRNA-encoding left and right TALENs designed to hit exon 3 of the *apoBa* gene (Supplemental Figure 1B) were injected into one-cell stage *Tg(fli1:eGFP)* embryos. We identified three different mutations, out of which we chose to focus on an 8-bp deletion in the TALENs target site, introducing a premature stop-codon after 70 amino acids (Supplemental Figure 1B). Both *apoBa* and *apoBb.1* mutants displayed strong decrease of the respective mRNA levels (Figure 1E). In addition, western blot analysis demonstrated complete absence of ApoB in *apoBb.1* but not in *apoBa* mutants (Supplemental Figure 1C), confirming previous data showing that ApoBb.1 is the predominant isoform (~95% abundance)^16^.

**Figure 1:**
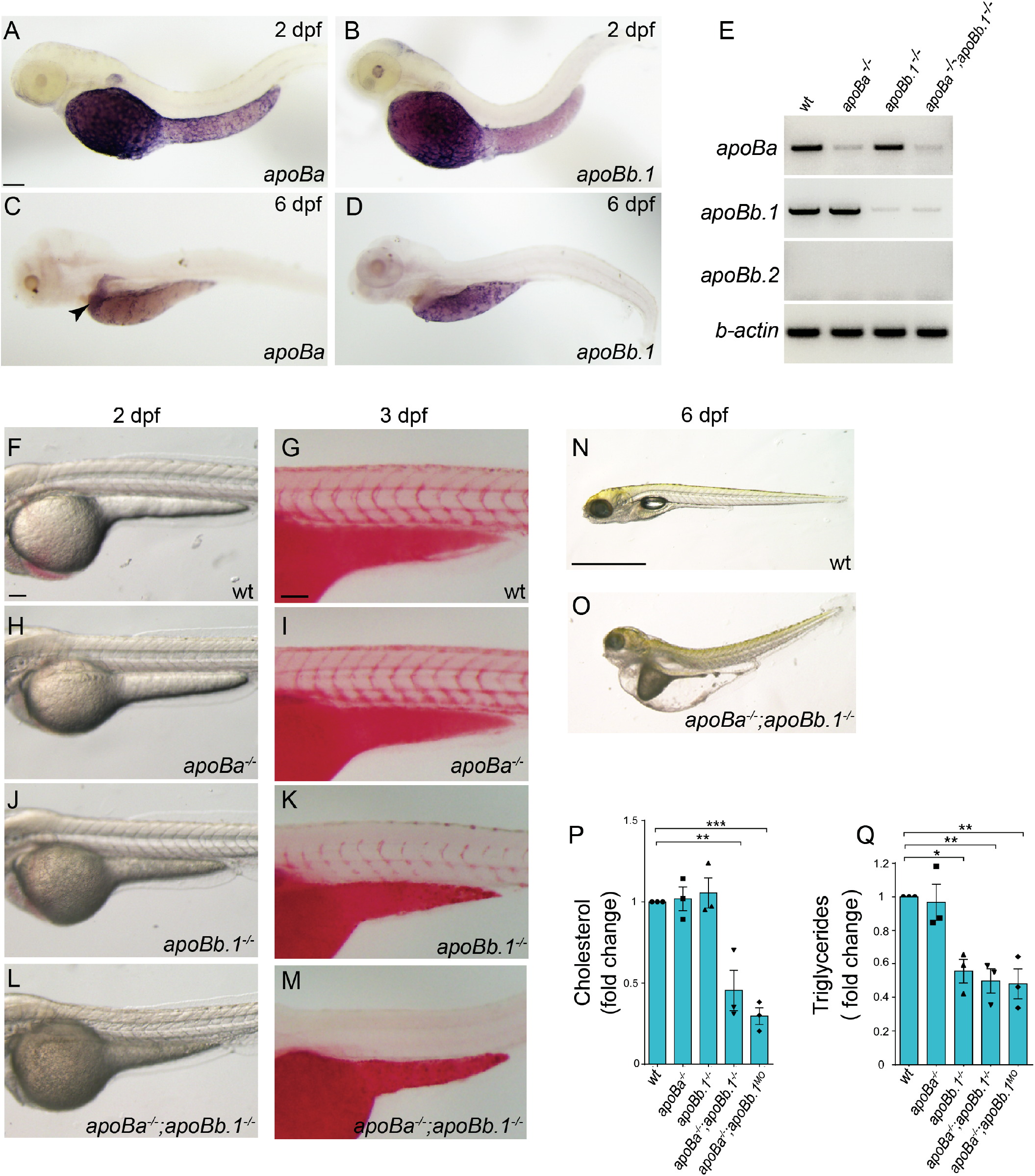
*apoB* mutants feature severe hypolipidemia. **(A-D)** WISH on 2 dpf zebrafish embryos showing strong expression of *apoBa* (A) and *apoBb.1* (B) in the YSL (n_*apoBa*_ =7; n_*apoBb.1*_=6). At 6 dpf *apoBa* expression is enriched in the liver (C) and *apoBb.1* in the YSL and the intestine (D) (n_*apoBa*_=3; n_*apoBb.1*_=3). **(E)** Semi-quantitative PCR for the *apoB* genes in the different mutants. **(F-M)** Transmitted light images of 2 dpf embryos showing dark yolk in *apoBb.1*^−/−^ (J), *apoBa*^−/−^;*apoBb.1*^−/−^ (L), but not in WT (F) or *apoBa*^−/−^ (H) mutant embryos. 3 dpf embryos stained with ORO show decreased lipid levels in *apoBb.1* mutant (K), as compared to WT (G) and *apoBa*^−/−^ (I) embryos. (M) *apoBa;apoBb.1* double mutants display complete absence of lipids in circulation. **(N,O)** Transmitted light images at 6 dpf demonstrate severe malformations, unabsorbed yolk and pronounced edema in *apoBa;apoBb.1* double mutants as compared to WT siblings. **(P,Q)** Cholesterol (P) and triglyceride (Q) levels in the different *apoB* mutants compared to WT controls at 3 dpf. N =3, n_embryos/sample_= 20. Scale bar: (A-D; F-M) 100 μm, (N,O) 1mm, P Value: *<0.05, **<0.01, ***<0.001.

### Characterization of the mutant phenotypes

*apoBa* homozygous mutants were viable, fertile and morphologically indistinguishable from their WT siblings (Figure 1F,H). Moreover, lipid distribution was normal in these mutants, as shown by Oil Red O (ORO) staining^17^ (Figure 1G,I). Similarly, injection of *apoBa* antisense morpholino oligonucleotides (MO) did not elicit any noticeable phenotypes (Supplemental Figure 2A,B). In contrast, *apoBb.1* homozygous mutants presented with several defects. First, they were easily identified due to their dark yolk (Figure 1J), indicative of impaired lipid absorption^17^. This was also verified by ORO staining, which showed a strong reduction in circulating lipid levels (Figure 1K). Injection of *apoBb.1* MO into wt embryos fully phenocopied the mutant phenotype (Supplemental Figure 2C), however no effects were observed following injection of *apoBb.1* MO into *apoBb.1* mutants (Supplemental Figure 4), confirming the MO specificity. The central role of the *apoBb.1* isoform was further corroborated by the decreased viability of the *apoBb.1* mutants (Supplemental Figure 2G). In addition, we noticed that the few *apoBb.1*^−/−^ animals that survived by ~60 dpf were significantly smaller than their WT counterparts (Supplemental Figure 2H).

**Figure 2:**
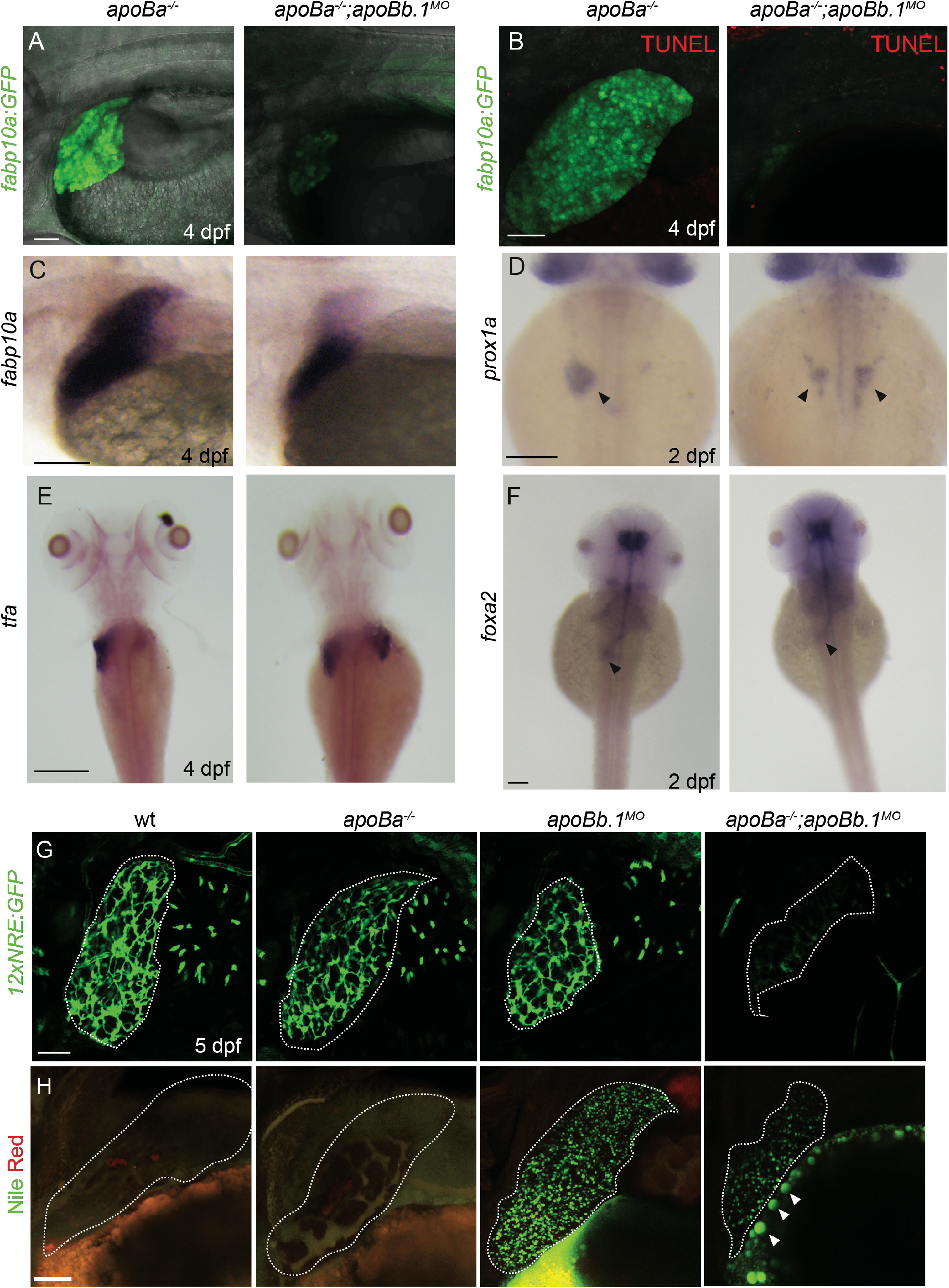
apoB mutants exhibit defective liver development and steatosis. **(A)** Confocal images at 4 dpf of *apoBa*^−/−^ and *apoba*^−/−^*;apoBb.1*^*MO*^ embryos on the background of *Tg(−2.8fabp10a:EGFP)*^*as3*^, showing smaller liver and decreased expression of *fabp10a EGFP* in *apoba*^−/−^*;apoBb.1^MO^* embryos (n_WT_=8/8, n_*apob*−/−;*apoBa*−/−*;apoBb.1MO*_=7/7). **(B)** No apoptotic cells are detected in the liver of *apoba*^−/−^ or *apoba*^−/−^*;apoBb.1*^*MO*^ embryos, following TUNEL staining (n_WT_=6/6, n_*apoBa*−/−;*apoBb.1MO*_=5/5). **(C)** WISH showing the spatial expression of *fabp10a* at 4 dpf, depicts decreased liver size in *apoba*^−/−^*;apoBb.1^MO^* embryo (n_WT_^-^=7, n_*apoBa*−/−;*apoBb.1MO*_=13). **(D,E)** WISH analysis with riboprobes against *prox1a* at 2 dpf (D) and *tfa* at 4 dpf (E) show bilateral liver buds in *apoba*^−/−^*;apoBb.1*^*MO*^ embryo as compared to *apoBa*^−/−^ mutants, which display a single liver bud located to the left of the midline. (**F**) *foxa2* expression at 2 dpf remains unchanged in *apoba*^−/−^*;apoBb.1*^*MO*^ embryos as compared to *apoba*^−/−^ siblings. **(G)** Confocal images of 5 dpf *Tg(12xNRE:Egfp)* embryos with outlined livers, showing decreased Notch activity in the bile ducts of *apoba*^−/−^*;apoBb.1*^*MO*^ embryos (n_WT_=4, n_*apoBa*−/−_=8, n_*apoBb*−/−_=6, n_*apoBa*−/−*;apoBb.1MO*_=20). **(H)** Confocal images of Nile Red staining at 4 dpf depicting neutral lipid accumulation in *apoBb.1*^*MO*^, and *apoba*^−/−^*;apoBb.1*^*MO*^ embryos. Red staining labels polar lipid, while green fluorescent highlights neutral lipids (n_WT_=5, n_*apoBa−/−*_=8, _n*apoBb.1−/−*_=5 n_*apoBa−/−;apoBb.1MO*_=9). Scale bar: (A,B,G,H) 50μm, (C-F) 100μm.

Although ApoBb.1 represents the predominant ApoB isoform, we suspected that the presence of ApoBa in *apoBb.1* mutants could compensate for the lack of its functions. Therefore, we generated *apoBa*;*apoBb.1* double homozygous mutants (Figure 1L). These animals displayed a darker yolk and complete absence of lipids in circulation (Figure 1M). Moreover, the lack of both ApoB isoforms led to lethality at 6-8 dpf, with the *larvae* exhibiting profound yolk and pericardial edema, as well as a curved trunk (Figure 1N,O). Finally, injection of *apoBb.1* MO into *apoBa* homozygous mutants or vice versa (*apoBa* MO into *apoBb.1*^−/−^ embryos) recapitulated the phenotypes of *apoB* double mutants (Supplemental Figure 1C and Supplemental Figure 2D-F).

Lipid profiling of the different mutants showed no differences in cholesterol levels between *apoBa*^−/−^ and *apoBb.1*^−/−^ individual mutants (Figure 1P). In contrast, these were strongly reduced in both *apoBa;apoBb.1* double mutants and *apoBa*^−/−^;*apoBb.1*^*MO*^, as compared to WT siblings (Figure 1P). Interestingly, triglyceride (TG) measurements revealed a slightly different picture (Figure 1Q). While in *apoBa*^−/−^ mutants TG levels were similar to those of WT siblings, they were strongly decreased in *apoBb.1* mutants (Figure 1Q). In addition, TG levels were significantly reduced in both *apoBa;apoBb.1* double mutants, and in *apoBa*^−/−^;*apoBb.1*^*MO*^ (Figure 1Q).

Due to the difficulty in obtaining large numbers of double homozygous mutants from *apoBa*^−/−^ x *apoBb.1*^+/−^ crosses, and having verified that the *apoBb.1* MO fully recapitulates the mutant phenotype, we decided to use *apoBa*^−/−^;*apoBb.1*^*MO*^ embryos in most experiments of this study.

### Defective organ development in *apoB* mutants

Next, we analyzed the development of the liver and the intestine, the main organs responsible for ApoB synthesis, assembly and secretion in vertebrates^2^. We took advantage of the fact that, in contrast to ApoB null mice, *apoBa*^−/−^;*apoBb.1*^−/−^ double mutants survive beyond the initial stages of formation of the digestive system, to investigate the role of ApoB during normal formation of these organs. To examine liver morphology, we used *Tg(−2.8fabp10a:EGFP)*^*as3*^, a well-established transgenic reporter expressing EGFP under the regulation of the *fatty acid-binding protein 10A (fabp10a)* promoter^23^. Injection of *apoBb.1* MO into *Tg(fab10a:EGFP);apoBa*^−/−^ embryos resulted in significantly reduced liver size at 4 dpf (Figure 2A and Supplemental Figure 3A), which was not due to apoptotic cell death, as confirmed by TUNEL staining (Figure 2B). Similar results were obtained following *in situ* hybridization for *fabp10a* (Figure 2C), suggesting a defect in hepatocyte specification, differentiation and/or proliferation. To distinguish between these possibilities, we analyzed the expression of *prox1a*, a transcription factor essential for hepatocyte differentiation and liver development^24,25^. As seen in Figure 2D, *prox1a*+ hepatic progenitors were normally detected to the left of the midline at 2 dpf, in phenotypically normal *apoBa*^−/−^ embryos. By contrast, loss of both *apoBa* and *apoBb.1* caused embryos to develop bilateral livers (Figure 2D, arrowheads), suggesting an early role for ApoB in controlling liver laterality. These defective positioning of hepatoblasts was still detected at 4 dpf, as depicted by the expression of *transferrin* (*tfa*), an additional liver-specific marker^26^ (Figure 2E). Interestingly, these defects were specific for the liver as we did not detect laterality problems in gut looping (Figure 2F, arrowheads and Supplemental Figure 3B) or in early left-right asymmetry markers such as *lefty1* (not shown). These results indicate that hepatoblast specification is initiated normally in *apoB* mutant embryos, but that liver formation begins to be affected after 2 days of development. Liver organogenesis in zebrafish involves two main phases: first, specification and migration of *hhex+* and *prox1+* hepatoblasts to form the liver bud on the left side of the embryo and second, hepatoblast differentiation into hepatocytes and biliary epithelial cells (BECs), followed by massive cell proliferation^27^. To check whether the development of the intrahepatic biliary network is also affected in *apoB* mutants, we used the *Tg(EPV.Tp1-Mmu.Hbb:EGFP)*^*ia12*^ reporter (12xNRE:EGFP), which consists of 12 repeats of Notch-responsive elements driving EGFP expression^28^, as Notch signaling in hepatic progenitor cells was shown to be both required and sufficient for biliary specification^29^. At 5 dpf, the *Tg(12xNRE:EGFP*) positive intrahepatic biliary network was highly branched in wt, *apoBa*^−/−^ and *apoBb.1*^−/−^ larvae (Figure 2G), as opposed to *apoBa*^−/−^;*apoBb.1*^*MO*^ that displayed largely reduced Notch activity. Taken together, the observed phenotypes suggest that, while ApoB is not required for initial hepatocyte specification, it appears to play an important role in hepatoblast positioning (i.e, budding and/or migration), proliferation and segregation between hepatocytes and BECs.

**Figure 3:**
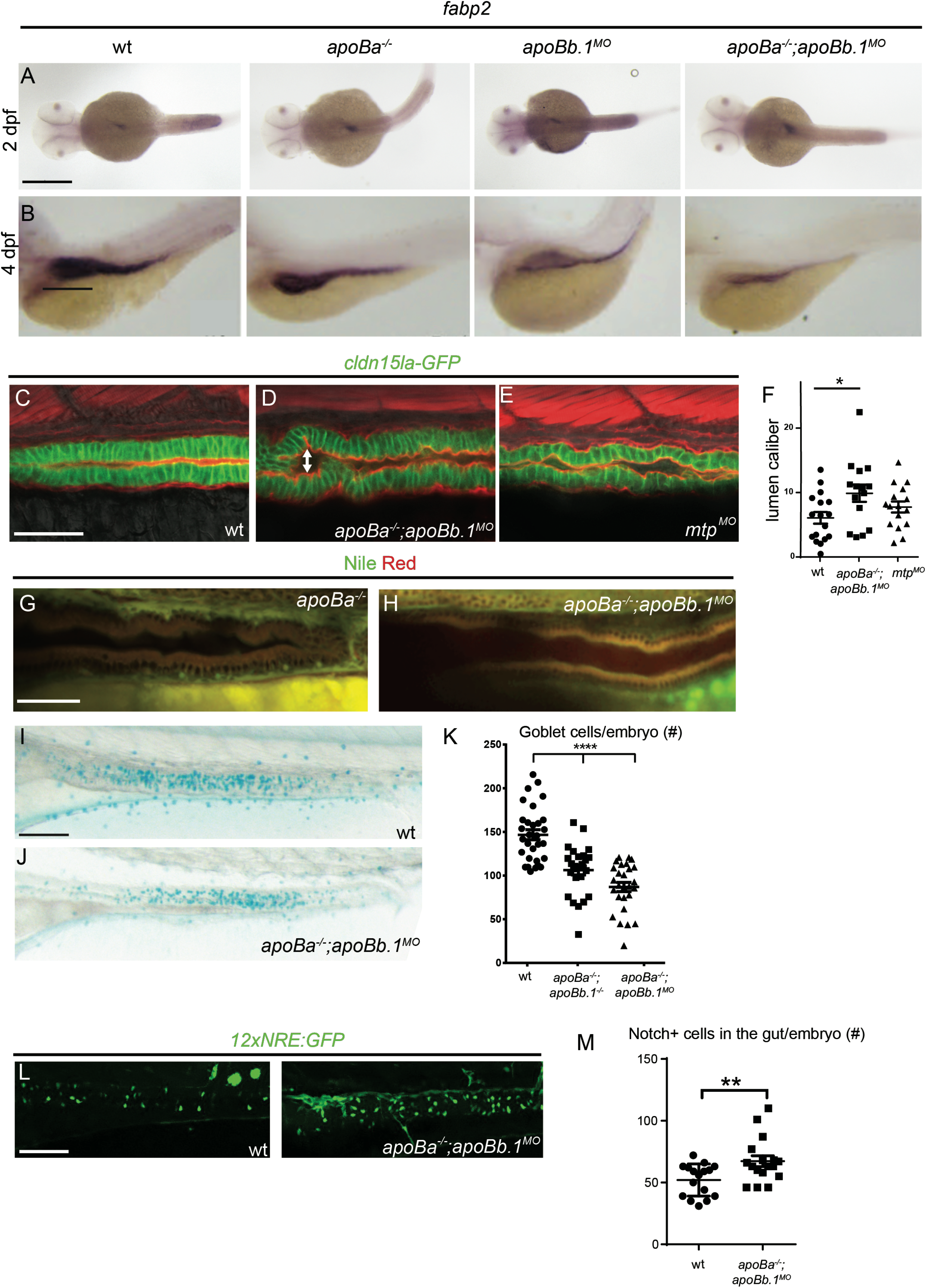
Gut development is impaired in *apoB* mutants. **(A,B)** WISH showing the spatial expression of *fabp2* at 2 (A) and 4 (B) dpf. **(C-E)** Confocal images at 4 dpf of wt (C), *apoba*^−/−^*;apoBb.1*^*MO*^ (D) and *mtp*^*MO*^ (E) embryos in the *TgBAC(cldn15la-GFP)*^*pd1034*^ reporter background, stained with Phalloidin (red). Enlarged lumen caliber (arrow) is detected in *apoba*^−/−^*;apoBb.1^MO^* and *mtp^MO^* embryos; quantified in (F) (N=2, n_wt_=17, n_*apoBa−/−;apoBb.1MO*_=15, n_*mtp MO*_=16). **(G,H)** Confocal images of the guts of 4 dpf *apoBa^−/−^* and *apoba*^−/−^*;apoBb.1*^*MO*^ embryos stained with Nile Red, show no lipid accumulation. **(I,J)** Bright field images of 5 dpf embryos stained with Alcian blue showing reduced number of goblet cells in *apoba*^−/−^*;apoBb.1*^−/−^ double mutants (J) as compared to wt embryos (I); quantified in (K) (N=3, n_WT_=30, n_*apoBa*−/−*;apoBb.1*−/−_=27, n_*apoBa−/−;apoBb.1MO*_=27). **(L)** Confocal images at 5 dpf of wt and *apoba*^−/−^*;apoBb.1*^*MO*^ embryos in the *Tg(12xNRE:Egfp)* reporter background. (M) Quantification of number of cells displaying active notch signaling in the guts of wt and *apoba*^−/−^*;apoBb.1*^*MO*^ embryos. Scale bar (A-E,G,H) 50μm, (I,J,L) 100 μm, P Value: *<0.05, **<0.01, ****<0.0001.

Hepatic steatosis, the accumulation of lipid within hepatocytes, is a critical step in the pathogenesis of several human diseases including alcoholic liver disease (ALD) and NAFLD. While NAFLD is mostly associated with hyperlipidemia^30^ defective synthesis and secretion of ApoB lipoproteins can also lead to hepatic steatosis, as observed in FHBL1 patients^9,31^. To investigate whether the different *apoB* mutants develop steatosis, we utilized Nile Red staining, which labels polar lipids in red and neutral lipids in green^32,33^. As expected, liver lipid stores were nearly undetected in WT and *apoBa*^−/−^ mutants (Figure 2H), as opposed to *apoBb.1*^*MO*^, which featured excessive accumulation of lipid droplets in hepatocytes (Figure 2H). Interestingly, lipid droplet accumulation was less severe in the livers of *apoBa*^−/−^; *apoBb.1*^*MO*^ embryos (Figure 2H), most probably due to the impaired secretion of ApoB lipoproteins, which remain stuck in the YSL (Figure 2H, arrowheads). These results were further confirmed by H&E staining of histological sections (Supplemental Figure 3C, arrowheads).

Because of the close relationship in the formation of the liver and the intestine we tested whether ApoB depletion affects also gut development. The intestine is responsible for the absorption of lipids, and their secretion in the form of chylomicrons into lymphatic vessels. Analysis of *foxa2*^34^ (Figure 2F, arrowheads and Supplemental Figure 3B) and *fabp2*^26^ (Figure 3A) expression revealed normal development of the intestinal tube in individual mutants at 2 dpf. However, by 5 dpf we noticed that the gut appears somewhat reduced in size in *apoBa*^−/−^; *apoBb.1* ^*MO*^ embryos (Figure 3B). Yet, the effects of ApoB depletion on gut formation appear to be less severe than the effects observed in the liver. We also inspected the guts of the different mutants by mating the fish with the *TgBAC(cldn15la-GFP)*^*pd1034*^ ^35^ reporter, which labels tight junctions in the intestine. We detected slightly enlarged lumens in the gut of *apoBa*^−/−^; *apoBb.1*^*MO*^ embryos (Figure 3C,D,F) as compared to WT siblings. Similar effects were observed following injection of *mtp* MO (Figure 3C,E,F). These defects were further corroborated by direct inspection of H&E staining of histological sections (Supplemental Figure 3D). While at this stage of development the intestine is known to be folded and to contain polarized epithelial cells with a microvillus brush border^36^ (Supplemental Figure 3D), in *apoBa*^−/−^; *apoBb.1*^−/−^ embryos, the intestinal epithelium appears thin, poorly folded and the cells have only few scattered microvilli (Supplemental Figure 3E). Interestingly, also intestine-specific deletion of *Mtp* or *ApoB* in mouse was shown to render small and large intestines with enlarged calibers^14,36^, mostly due to fat accumulation in the villus enterocytes, fully recapitulating the phenotypes reported for abetalipoproteinemia^37^. In contrast, we did not detect lipid accumulation in enterocytes of *apoBa*^−/−^; *apoBb.1*^−/−^ mutants (Figure 3G,H), suggesting that the enlarged lumen phenotype and thinner epithelia are most likely derived from impaired nutrient absorption, as previously described^38^.

Finally, Alcian blue staining at 5 dpf highlighted a significantly reduced number of goblet cells in *apoBa*^−/−^; *apoBb.1*^−/−^ mutant guts (Figure 3I-K). Previous reports have shown that inhibition of Notch signaling directs the specification of intestinal cells toward a secretory fate^39^, leading to increased numbers of goblet cells. We therefore evaluated the levels of Notch activation in the guts of wt and mutant embryos using the 12NRE:GFP reporter. As seen in Figure 3L,M, we detected increased numbers of Notch+ cells in the guts of *apoBa*^−/−^; *apoBb.1*^*MO*^ embryos as compared to wt siblings, pointing to alteration in Notch signaling as the mechanism underlying goblet cell differentiation. Altogether, our results indicate that ApoB deficiency results in impaired liver development accompanied by hepatic steatosis and disrupted intestinal architecture.

### ApoB depletion affects vascular development

Previous studies, including our own have demonstrated that ApoB-containing lipoproteins have an inhibitory effect on vascular growth^11,17,18,40^. We therefore decided to analyze the involvement of the ApoB isoforms in the development of the vascular system by assessing the early vasculature of the different mutants. We first focused on the subintestinal vessels (SIVs), a plexus located on top of the yolk syncytial layer (YSL) (Figure 4A) that plays a critical role in yolk absorption and vascularization of the gastro-intestinal tract^17,28,41^. In line with the lipid and morphological defects, the vascular phenotypes were more pronounced in the double homozygous mutants, as compared to WT siblings or single mutants. As seen in Figure 4B-D,H-I, the SIVs formed normally in *apoBa*^−/−^ and displayed only minor defects in *apoBb.1* mutants. By contrast, *apoBa*;*apoBb.1* double mutants (Figure 4E,H,I) and *apoBa*^−/−^;*apoBb.1*^*MO*^ (Figure 4F,H,O) larvae exhibited ectopic angiogenic sprouts arising in the SIVs (Figure 4G)^17^, which did not retract during the remodeling phase, as in wt siblings^28^. These sprouts, reminiscent of those observed in *stl* mutants were clearly detected following injection of *apoBb.1* MO into wt embryos (Figure 4E,F), however no effects were observed upon injection of *apoBb.1* MO into *apoBb.1* mutants (Supplemental Figure 4B-E), confirming the MO specificity. Similar hyperangiogenic behaviors were detected in the posterior cerebral vein (PCeV) at 4 dpf (Supplemental Figure 4A,F-H), and in the dorsal trunk (Supplemental Figure 4A,I-K) of *apoBa*^−/−^;*apoBb.1*^*MO*^ larvae, suggesting that impaired angiogenesis is a general consequence of ApoB depletion, which becomes more pronounced as development proceeds.

**Figure 4:**
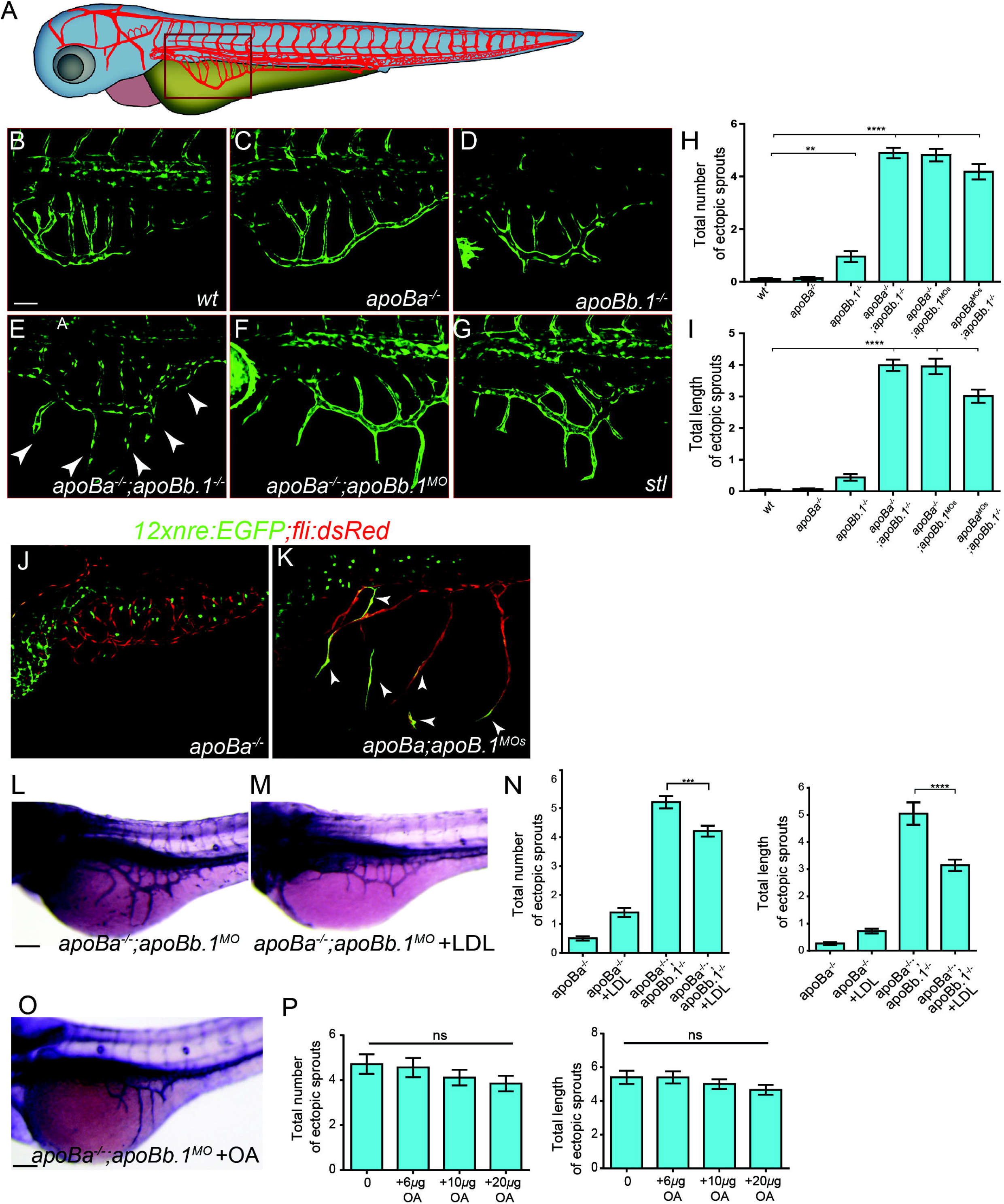
apoB mutants display hyperangiogenic phenotypes. **(A)** Schematic representation of the zebrafish embryonic vasculature, with red square marking the subintestinal vessels (SIVs). **(B-G)** Confocal images at 3 dpf showing ectopic sprouts (arrowheads) arising in the SIVs of *apoba*^−/−^*;apoBb.1*^−/−^ double mutants (E), *apoba*^−/−^*;apoBb.1*^*MO*^ (F) and *stl* mutants (G), but not in WT (B), *apoBa*^−/−^ (C) or *apoBb.1*^−/−^ (D) individual mutants. **(H,I)** Quantification of the number (H) and length (I) of ectopic sprouts at 3 dpf. N=3, n_WT_=45, n_*apoBa*−/−_=55, n_*apoBb.1−/−*_=23, n_*apoBa*−/−*;apoBb.1*−/−_=31, n_*apoBa*−/−*;apoBb.1MO*_=40, n_*apoBb.1*−/−*;apoBaMO*_=21. **(J,K)** Confocal images at 5 dpf of wt and *apoba*^−/−^*;apoBb.1*^*MO*^ embryos in the *Tg(12xNRE:Egfp)* reporter background. White arrowheads point to ECs with active Notch signaling in ectopic sprouts that failed to retract in *apoBa*^−/−^;*apoBb.1*^*MO*^ embryos. **(L,M)** Transmitted light images of the SIVs at 3 dpf, showing inhibition of ectopic sprouting following intravascular injection of DiI-LDL into *apoBa*^−/−^*;apoBb.1*^*MO*^. (N) Quantification of number and length of ectopic sprouts following intravascular injection of DiI-LDL (N=3, n_*apoBa*−/−_=57, n_*apoBa*−/−+LDL_=43, n_*apoBa*−/−*;apoBb.1MO*_=67, n_*apoBa*−/−*;apoBb.1MO*+LDL_=57). **(O)** Transmitted light images of the SIVs of 3 dpf *apoBa*^−/−^;*apoBb.1*^*MO*^ embryos treated with OA, quantified in (P) (N=3, n_*apoBa*−/−*;apoBb.1MO*_=20, n_*apoBa*−/−*;apoBb.1MO* + 6*μg OA*_=19, n_*apoBa*−/−*;apoBb.1MO*+*10μg OA*_=21, n_*apoBa−/−;apoBb.1MO+20μg OA*_=22). Scale bar (B-G) 50μm, (J-O) 100 μm, P Value: ****<0.0001, ***<0.001, **<0.01.

We have previously shown that ApoB-lipoproteins induce ectopic angiogenesis by downregulating the expression of the VEGF-decoy receptor Vegfr1/Flt1^17^. Accordingly, depletion of both membranous and soluble forms of Flt1 (mFlt1 and sFlt1 respectively) fully phenocopied this phenotype^28^. Given that loss of Flt1 has been shown to result in enhanced Notch signaling^42^, and that Flt1^−/−^ mutant sprouts are less likely to retract^43^, we decided to investigate whether this mechanism is activated in *apoB* mutants. Live imaging of the *12xNRE:EGFP* reporter revealed clear activation of Notch signaling in the ectopic sprouts that failed to retract in *apoBa*^−/−^;*apoBb.1*^*MO*^ embryos (Figure 4J,K). In contrast, no ECs displaying active Notch signaling were detected in the SIV plexus of wt embryos at comparable developmental stages (i.e., 5 dpf), when the vessels have acquired their final, quiescent stage. Taken together these results reinforce and expand our previous findings by showing that not only Flt1, but also Notch signaling act downstream to ApoB to modulate angiogenesis.

In order to verify that the observed vascular phenotypes indeed derive from the absence of ApoB lipoproteins, we attempted to rescue the defects by intravascularly injecting DiI-labeled LDL into *apoBa*^−/−^;*apoBb.1*^*MO*^ embryos (Supplemental Figure 4L) as previously described^17,19^. As seen in Figure 4L-N, a single injection of DiI-LDL was sufficient to revert the excessive angiogenesis phenotype, as opposed to oleic acid (OA) supply (Figure 4O,P), supporting a specific role for ApoB, and not the lipid moieties within lipoproteins, in developmental angiogenesis. These results are in line with our previous findings, which showed that the strong excessive angiogenesis phenotype displayed by *stl* mutants in response to lipoprotein depletion (Figure 4G) was efficiently reversed following injection of a delipidated form of ApoB^17^.

Because this pathway could potentially be exploited to control pathological neovascularization, we proceeded to define the domains within ApoB that act on ECs. To this end we tested the ability of truncated forms of the human APOB protein to inhibit angiogenesis, using the well-established tube formation assay. To ensure that lipoproteins are presented to the cells in their physiological form, we co-transfected MTP along with human APOB34 or APOB25, which have been previously shown to undergo proper lipidation and secretion^44,45^, into HEK cells (Supplemental Figure 4M). We then transferred the media containing the different truncated forms to HUVEC cells, and examined their ability to generate tubes. LDL, which we previously showed inhibits tube formation in culture^18^, was used as positive control. Interestingly, both APOB25 and APOB34 forms inhibited angiogenesis in this assay, suggesting that the anti-angiogenic effect of ApoB is conveyed by the 25% (N-terminus) of the protein (Figure 5A-E).

**Figure 5:**
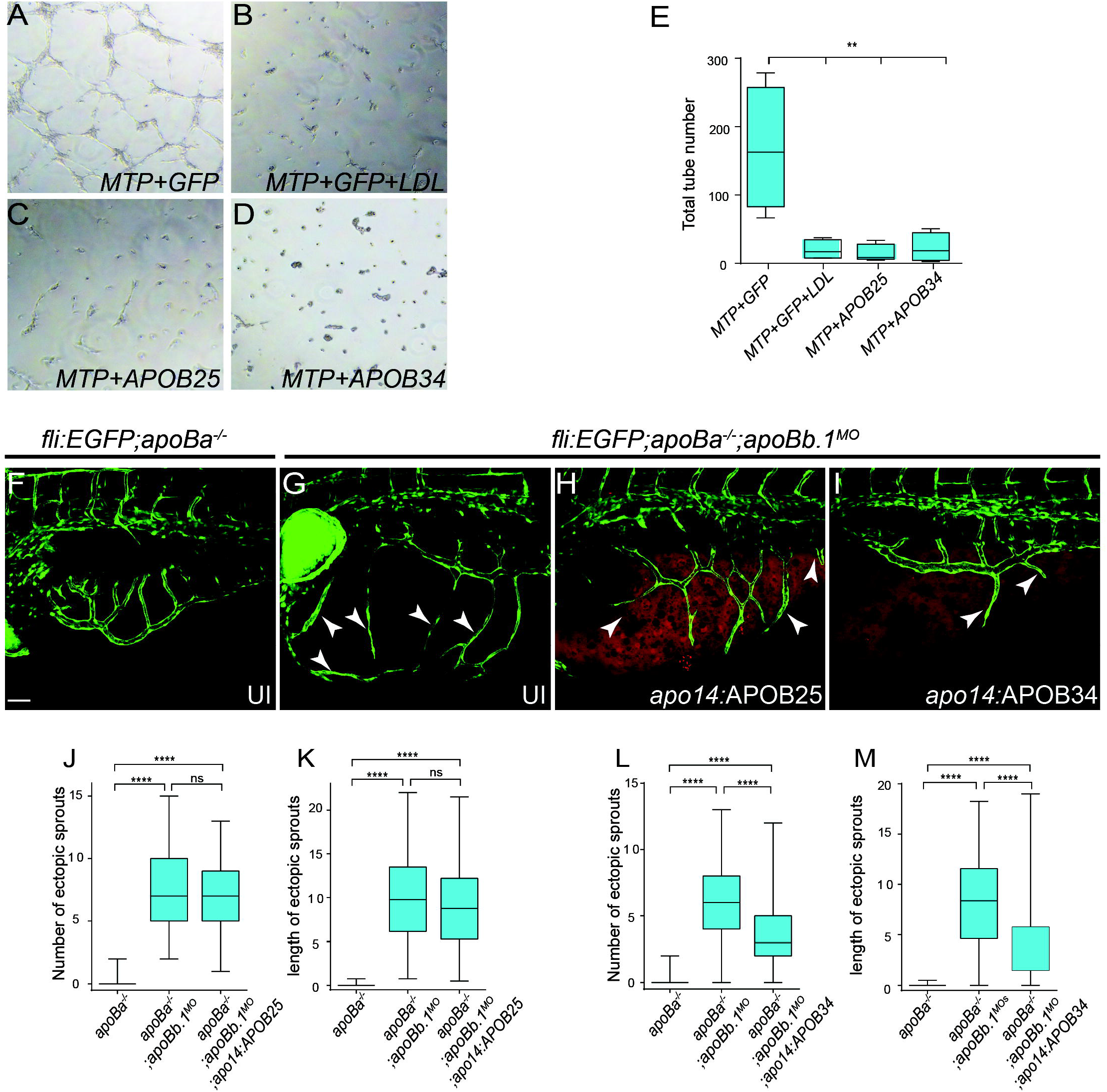
Truncated forms of human APOB inhibit angiogenesis in *apoB* mutants. **(A-D)** Tube formation assay on HUVECs untreated (A,C,D) or treated with LDL (B) plus conditioned medium from HEK293 cells co-transfected with (A,B) *MTP+GFP,* (C) *MTP+APOB25* and (D) *MTP+APOB34*. **(E)** Quantification of total number of tubes following different treatments. N=3, n_*MTP*+*GFP*_=4, n_*MTP*+*GFP*+*LDL*_=4, n_*MTP*+*APOB25*_=5, n_*MTP*+*APOB34*_=4 **(F-I)** Confocal images at 4 dpf showing reduction in the number and length of ectopic sprouts in *Tg(fli1:eGFP);apoBa*^−/−^*;apoBb.1*^*MO*^ embryos injected with *apo14:APOB34* (I) as compared to uninjected *apoBa*^−/−^ (F), uninjected *apoBa*^−/−^;*apoBb.1*^*MO*^ (G), and *apoBa*^−/−^;*apoBb.1*^*MO*^ injected with *apo14:APOB25* (H). **(J-M)** Quantification of the number and length of ectopic sprouts in *apo14;APOB25* (J,K) (N =3, n_*apoBa*−/−_=47, n_*apoBa*−/−*;apoBb.1MO*_=54, n_*apoBa−/−;apoBb.1MO;apo14:APOB34*_=54), and *apo14;APOB34* (L,M) (N=3, n_*apoBa−/−*_=47, n_*apoBa−/−;apoBb.1MO=*_51, n_*apoBa*−/−*;apoBb.1MO;apo14:APOB34*_=55) injected embryos. Scale bar (F-I) 50μm, P Value ****<0.0001, **<0.01

We then asked whether these fragments are sufficient to inhibit angiogenesis *in vivo*. To answer this question we generated plasmids expressing human *APOB25* and *APOB34* fused to *IRESmCherry*, under the regulation of the *apo14* promoter, which drives expression in the YSL and the liver^46^. While the hyperangiogenic phenotype of *apoBa*^−/−^;*apoBb.1*^*MO*^ embryos was not reverted following *apo14:APOB25-IRESmCherry* injection (Figure 5G,H,J,K), *apoBa*^−/−^;*apoBb.1*^*MO*^ embryos injected with *apo14:APOB34-IRESmCherry* displayed a significant reduction of both the number and length of the ectopic sprouts at 4 dpf (Figure 5G,I,L,M). These results suggest that the first 34% of the ApoB protein are required to exert its anti-angiogenic effect *in vivo*. The discrepancy between the *in-vivo* and the *in-vitro* results may derive from differences in the mobility and clearance of the truncated forms of APOB *in vivo*. Specifically, it has been shown that APOB25 is not present in circulation *in vivo*, despite of it being produced and secreted to the blood^44,47^, due to fast clearance.

As a whole these results demonstrate the applicability of our newly generated models of ApoB deficiency for assessing the physiological activity of potential therapeutic agents.

## Discussion

In this work, we generated zebrafish mutants for two ApoB isoforms: *apoBa* and *apoBb.1*. Similar to null *ApoB* mice, zebrafish carrying mutations in both *apoBa* and *apoBb.1* result in early lethality. However, in contrast to the mammalian models, the external fertilization and development of zebrafish allowed us to investigate previously unappreciated roles of ApoB during embryogenesis.

Three *apoB* genes are present in the zebrafish genome, namely *apoBa*, *apoBb.1* and *apoBb.2*. During early development, *apoBa* and *apoBb.1* are highly expressed, whereas *apoBb.2* is absent^16^. Interestingly, despite the strong expression of *apoBa* mRNA, its protein levels are almost twenty-fold lower than those of ApoBb.1^16^. Accordingly, while the absence of *apoBa* did not elicit any noticeable phenotype, *apoBb.1* depletion caused strong hypolipidemia. Finally, deletion of both isoforms resulted in severe phenotypes and lethality.

Our results demonstrate novel roles for ApoB during development the liver, one of its main producing organs. In particular, we find that *apoBa*^−/−^;*apoBb.1*^*MO*^ embryos display similar phenotypes as those caused by retinoic acid (RA) deficiency^48^, included decreased liver volumes and liver bud bilaterality. Zebrafish embryos treated with the Raldh inhibitor DEAB, or injected with MOs against the three RA receptors (*rarab*, *raraa*, and *rarga)* were shown to display smaller livers as a result of impaired hepatocyte proliferation. In contrast, RA addition to the fish water resulted in massive proliferation of hepatocytes and enlarged livers^48^. In addition, downregulation of the RA receptor *rargb* led to bilateral liver buds and intrahepatic biliary defects, similar to those displayed by double *apoB* mutants. While most of RA circulation is mediated through retinol binding protein 4 (RBP4), it has been shown that ApoB-containing lipoproteins (e.g. LDL and vLDL) also participate in RA shuttling^49^. Therefore, the possibility exists that complete depletion of ApoB leads to RA unavailability, which in turn results in liver bilaterality and reduced hepatocyte and BEC proliferation. The absence of both apoB isoforms resulted in low levels of triglycerides and cholesterol in circulation, accompanied by massive lipid droplet accumulation in hepatocytes, and liver steatosis. These phenotypes, reminiscent of human NAFLD, recapitulate part of the clinical manifestations of FHBL1, making the *apoBa*^−/−^;*apoBb.1*^−/−^ mutant zebrafish an advantageous model for the study of this disorder and for the identification of potential treatments.

In addition to liver abnormalities, ApoB and MTP deficiency resulted in disrupted intestinal architecture. In particular, the guts were poorly folded and displayed enlarged lumen calibers, resembling phenotypes displayed by starved wt larvae^38^. Double *apoB* mutants feature also increased number of goblet cells. Interestingly, it has been shown that retinoic acid negatively regulates goblet cell differentiation^50^, whereas inhibition of Notch signaling directs specification of intestinal cells toward a secretory fate^39^ leading to increased numbers of goblet cells. Our results indeed demonstrate increased activation of Notch signaling in the intestines of *apoBa*^−/−^;*apoBb.1*^−/−^ mutants, supporting the idea that ApoB affects goblet cell differentiation through activation of Notch targets. Downstream to Notch, the transcription factor Klf4, has been shown to be required for terminal differentiation of goblet cells^51^. *klf4a* expression was shown to be controlled by Notch signaling (i.e., embryos treated with the γ-secretase inhibitor DAPT display increased *klf4a* expression in the intestine, while decreased *klf4a* expression and reduction in goblet cell number were observed in embryos injected with *Notch intracellular domain* (*NICD*) mRNA^52^). On the other hand, it has been demonstrated that the RA-RARa axis negatively modulates *klf4* expression, thereby impacting as well goblet cell differentiation. In the future it will be interesting to investigate whether ApoB is involved in the interplay between the Notch and RA-RARa pathways in the developing intestine. Finally, the phenotypes detected in the liver and intestine of *apoB* mutants could potentially result from the absence of fat-soluble vitamins, such as vitamins E and A, and their Retinoic acid (RA) metabolites, whose absorption and transport throughout the body rely mostly on ApoB, in the form of chylomicrons^53,54^.

Previous studies have demonstrated the strong impact of lipoproteins on endothelial cell behavior^17,19,55,56^; yet, the mechanisms by which ApoB affects these cells is not fully understood. The double *apoB* mutant exhibited a marked angiogenic phenotype characterized by excessive sprouts in the SIV, PeCV and in the aISVs. This angiogenic effect may result from a combination of different factors related to ApoB deficiency, such as lipid raft density, lack of fat-soluble vitamins and activation of downstream signaling pathways in ECs. Interestingly, the angiogenic defects were observed only following complete depletion of circulating ApoB-lipoproteins. Although *apoBb*.*1* mutants featured severe hypolipidemia, they did not display a strong angiogenic phenotype, suggesting that small amounts of ApoB are sufficient to maintain healthy control over the angiogenic process and further confirming the specific effects of the ApoB protein on EC behavior. Our structure-function analyses revealed that APOB34, a truncated form of human APOB lacking the LDL receptor binding domain, is able to rescue the angiogenic phenotype in double *apoB* mutants. These results raise the provoking hypothesis that an LDLR-independent pathway might regulate the effects of ApoB on ECs. Recently, it has been shown that LDL can bind the TGF-β receptor activin receptor-like kinase 1 (ALK1)^57^, which is also involved in lipoproteins transcytosis^58^. Moreover, ALK1 has been shown to exert anti-angiogenic effects via up-regulation of VEGFR1^59^. Interestingly, *stl* mutants display reduced expression of *vegfr1*, leading to excessive angiogenesis^17^. Therefore, it seems feasible that APOB induction of ALK1 activity leads to up-regulation of *vegfr1* and inhibition of angiogenesis.

Overall, our newly generated *apoB* mutants can serve as a model for studying human pathologies associated with hypolipidemia, as well as for related drug screens. Moreover, understanding the molecular mechanisms by which ApoB regulates angiogenesis will provide new therapeutic targets for the treatment of vascular pathologies ranging from cancer to ischemic heart disease.

## MATERIALS and METHODS

### Zebrafish husbandry and transgenic lines

Zebrafish were raised by standard methods and handled according to the Weizmann Institute Animal Care and Use Committee. *Tg(fli1:EGFP)*^*yl*^, *stl*^17^, *TgBAC(cldn15la-GFP)*^*pd1034*^ ^35^, *Tg(fli1:dsRed)*^*um13*^ ^60^, T*g(lyve1:dsRed2)*^*nz101*^ ^61^, *Tg(flt1_9a_cFos:GFP)*^*wz2*^ ^61^, *Tg(EPV.Tp1-Mmu.Hbb:EGFP)*^*ia12*^ ^28^ and *Tg(−2.8fabp10a:EGFP)*^*as3*^ ^23^ have been previously described. To generate *apo14:APOB25-IRESmCherry* and *apo14:APOB34-IRESmCherry*, the zebrafish *apo14* promoter^46^ was cloned into pDONRP4-P1R (Invitrogen), human *APOB25* and *APOB34* ^44^ were cloned into pDONR221 (Invitrogen), and the *IRESmCherry* sequence was cloned into pDONRP2R-P3 (Invitrogen), using Gateway BP Clonase II (Invitrogen, 11789-020). The three vectors were then transferred into pDestTol2pA2^60^ using a Gateway LR reaction (Invitrogen, 12538-120) The plasmids were injected along with Tol2 transposase mRNA into one-cell-stage embryos^60^.

### Generation of zebrafish mutants

1. *apoBb.1* CRISPR guide was designed with CHOPCHOP (chopchop.rc.fas.harvard.edu), potential off-target sequences were assessed using the MIT CRISPR Design site (crispr.mit.edu). Oligonucleotides synthesized for the guide sequence GGACTAGTGTGGTCTTTGAC were cloned into the BsmBI sites of the pT7-gRNA plasmid (Addgene, 46759). Cas9 mRNA was generated from pCS2-nCas9n^62^ using mMACHINE T7 ULTRA kit (Ambion, AM1345). Cas9 mRNA (250 ng/μl) and gRNAs (100 ng/μl) were co-injected into one-cell-stage *Tg(fli1:EGFP)*^*y1*^ transgenic embryos. For genotyping, genomic DNA was extracted at 24 hpf, amplified with 5’-TACTCTGTGAATGCCAGACAGG and 5’-TCAGAAGCTATCCAGCAAACA (184bp) primers, and analyzed on 3% Agarose gel.
2. *apoBa* TALENs left - TACAGCTAACCTCAAGAA and right - TGCCAGgtacaaaaaca plasmids were generated as described^63^ and transcribed using mMESSAGE mMACHINE T7 ULTRA kit (Ambion, AM1345). mRNAs (250ng each) were co-injected into one-cell-stage. For genotyping, genomic DNA was extracted at 24 hpf, amplified with 5’-TATCAGTACACAGCAGAGAGCA and 5’-TCGTAAAATTAGGCTAAGCCA (110bp) and analyzed on 3% Agarose gel.

### Antisense morpholino injection

The following antisense morpholino oligonucleotides (Gene-tools) were re-suspended and injected as described^64^ at the following concentrations: *mtp* (4 ng), *apoBa* (4ng), *apoBb.1* (2ng).

### *In situ* hybridization and staining procedures

Embryos were fixed overnight in 4%PFA and processed for Oil Red O (ORO)^17^, Alkaline Phosphatase (AP)^17^, Nile Red (sigma, N3013)^33,65^, and Alcian Blue (Sigma, A3157)^66^ staining, as described. TUNEL was performed using *In Situ* Cell Death Detection Kit, TMR red (Roche, 12156792910) following manufacturer’s instructions.

*In situ* hybridization was performed as described^67^ using the following probes: *fabp10a*^68^, *prox1a*^69^,. The following primers were used to generate the corresponding riboprobes:

*apoBa* 5’-GACCTTGGCTTTCCGTTCC-3’, 5’-CAGCAGGGAAGCTCTCTATGAA-3’,

*apoBb.1* 5’-GCTGCAGTGTATGCCATGGGAAT-3’, 5’-ATCAACAGTGGGTTCCAGACCCTT-3’.

*foxa2* 5’-GTGTTACACCTCGGTCAGCA-3’, 5’-CACTTGAAGCGCTTTTGCCT-3’

*fabp2* 5’-TGGAAAGTCGACCGCAATGAGA-3’, 5’-TACCTTTCCGTTGTCCTTGCGT-3’

*tfa* 5’-TCTGGAGGCTGGAATACTCCTA-3’, 5’-TAAACCTGAGCCCTTACGCA-3’

### Assessment of Vascular Phenotypes

*Larvae* were assayed for SIV ectopic sprouting using fluorescent imaging (PCeV, Trunk) or AP staining (SIVs). Quantitation of the SIVs phenotype was done using a grid lens in a Leica Stereoscope under X10 magnification, each side of the animal was scored separately.

### Total RNA isolation and semi-quantitative RT-PCR

A pool of 10-20 embryos/sample was homogenized in Trizol (Invitrogen, 15596026) and processed for RNA extraction following standard procedures^17^. 1μg of total RNA per reaction was reverse transcribed using a High-Capacity cDNA Reverse Transcription Kit (Applied Biosystem, 4368814). To measure relative changes in mRNA transcripts we used the following primers:

*apoBa* 5’-GACCTTGGCTTTCCGTTCC-3’, 5’-CAGCAGGGAAGCTCTCTATGAA-3’,

*apoBb.1* 5’-GCTGCAGTGTATGCCATGGGAAT-3’, 5’-ATCAACAGTGGGTTCCAGACCCTT-3’

*apoBb.2* 5’-GTTCATAGGAGCGAGCATTGACCA-3’, 5’-AGACCCAAACTGTCAACGAAAGGC-3’.

Expression levels were standardized to the primer set specific for β-actin 5’ TGACAGGATGCAGAAGGAGA-3’ - and 5’-GCCTCCGATCCAGACAGAGT -3’

### Triglyceride and cholesterol measurements

20 deyolked embryos at 3dpf per sample were gently homogenized in PBS with a pestle. After centrifugation at 15,000 g for 15 min, supernatants were collected and triglyceride and cholesterol levels were measured using kits from BioVision (Milpitas, CA; Triglyceride Quantification Kit, K622-100; Cholesterol Quantification Kit, K623-100) as described^70^. For each genotype, 3 repeats of 10-20 embryos each, were performed.

### Protein extraction and Western blot

3 dpf zebrafish embryos were processed for Western Blot as described^69^. Briefly, proteins were separated by SDS-PAGE using a 6% separating gel and a 4% stacking gel. Transfer was performed at 400 mA for 1h. Membranes were blocked with 2% BSA in PBST. The following antibodies were used:

Rabbit anti-human APOB100 (Abcam, ab20737), 1:1500.

Mouse monoclonal anti-human αTubulin (Sigma, T5168), 1:3000.

### Oleic acid and LDL treatments

Oleic acid (Sigma, O7501) was dissolved in ethanol and added to the fish water with fatty acid free BSA (SIGMA, A8806), at 6-20 μg/ml concentration.

Dil-LDL (Invitrogen, L3482) was injected intravascularly at 2 dpf, as described^17^.

### Imaging

Confocal imaging was performed using a Zeiss LSM 780 upright confocal microscope (Carl Zeiss, Jena, Germany) with a W-Plan Apochromat ×20 objective, NA 1.0. Images were processed using ImageJ (NIH). Fluorescent proteins were excited sequentially with 488 nm and 563 nm single-photon lasers.

### Cell transfection and culture

50% confluent HEK293 cells were co-transfected with 5ug *Huh7MTP* ^17^ and 10ug pGFP or *APOB34-FLAG* or *APOB25-DsRed*^44^, using JET PEI (Polyplus, catalog no.15021C1T) standard protocol. 24h after transfection, medium was replaced with EBM, containing 0.5% LPDS and 1% penicillin-streptomycin. 48 hrs later the medium was collected and centrifuged at 2000 rpm for 5min to ensure a cell free media. The upper phase was used as conditional medium for the tube formation assay.

### Tube formation assay

HUVECs (Lonza) were cultured on gelatin coated dishes, in M199 medium supplemented with 20% FCS, 50μg/ml Endothelial Cell Growth Supplement (ECGS, Zotal catalog no.BT-203), 5units/ml heparin, 1% pen-strep, 2mM L-glutamine 2mM^18^.

For treatment, conditional medium collected from *MTP*+*GFP*, *MTP*+*GFP*+LDL (100μg/ml) (BT-903), *MTP+APOB25* and *MTP+APOB34* transfected HEK293 cells, was added. Following 18h incubation, 30,000 cells were seeded on Matrigel© (BD, 356231) for 8 hrs, and cultured with the appropiate conditional medium (*MTP*+GFP, *MTP+*GFP+LDL, *MTP*+*APOB25* and *MTP*+*APOB34*). Cells were then fixed in 4% PFA and imaged using bright-field microscope. 9 fields/well were acquired per experiment. Total tube numbers were quantified using imageJ. For each experiment, values were normalized to the non-treated well.

### Statistical Analyses

All data are reported as mean values ± SEM and were analyzed using Prism 5 software (GraphPad Software, Incorporated, La Jolla, CA, USA). Comparison of two samples was done by unpaired two-side student *t*-test. Statistical significance for three or more samples was calculated via one-way ANOVA followed by post hoc Tukey’s for multiple comparisons. In box&whisker graph, box- 25th to 75th percentiles, line-median, whisker represents min and max.

## Author contributions

H.T. designed and conducted all experiments, analyzed data, and co-wrote the manuscript; I.A-D. conducted HUVEC experiments; N.M. managed fish work and conducted ZF experiments; K.Y. initiated and directed the study, designed experiments, analyzed data and co-wrote the paper with input from all authors.

## Acknowledgments

The authors would like to thank Yona Eli, Hila Raviv and Lital Shen (Weizmann Institute, Israel) for technical assistance, S. Ben-Dor (Weizmann Institute, Israel) for bioinformatic analysis, Noa Stettner, Alon Harmelin, Gabriella Almog, Roy Hofi, and Anna Tatarin for superb animal care, Didier Stainier (Max Planck Institute for Heart and Lung Research, Germany) for providing *Tg(−2.8fabp10a:EGFP)*^*as3*^ fish, Michel Bagnat (Duke University Medical Center, NC) for providing the *TgBAC(cldn15la-GFP)*^*pd1034*^ transgenic line, GregoryS. Shelness (National Institutes of Health, MD) for the APO25and APOB34 plasmids. The authors are grateful to all the members of the Yaniv lab for discussion, technical assistance and continuous support. This work was supported in part by European Research Council 335605 (to K.Y.), Israel Science Foundation 861/2013 (to K.Y.), the H&M Kimmel Inst. for Stem Cell Research, and the Estate of Emile Mimran (SABRA program), the Willner Family Center for Vascular Biology; the estate of Paul Ourieff; the Carolito Stiftung; Lois Rosen, Los Angeles, CA; Edith Frumin; the Fondazione Henry Krenter; the Wallach Hanna & Georges Lustgarten Fund, the Polen Charitable Trust and the Daniel Shapiro Cardiovascular Fund. K.Y. is the incumbent of the Enid Barden and Aaron J. Jade Professorial Chair in Memory of Canter John Y. Jade

---

**Supplemental Figure 1: Generation of *ApoB zebrafish mutants.* (A)** Schematic representation of the *apoBb.1* genomic locus, with the guide target site in exon 5. Guide recognition site is shown in green and the PAM in red. The mutant sequence carries an 18bp insertion containing an in frame stop-codon (TAG). **(B)** *apoBa* genomic locus, with the putative TALEN site in exon 3. The sequence contains TALENs recognition sites (blue), and a TALEN spacer (16 nucleotides, green) with the 8-bp deletion in the mutant. **(C)** Western blot analysis showing complete depletion of the ApoB protein in the *apoBb.1* mutant and in double mutants.

**Supplemental Figure 2**: **(A-C)** 3 dpf embryos injected with *apoBb.1* MO (C) show decreased lipid levels, as compared to embryos injected with control MO (A) OR *apoBa* MO (B). **(D-F)** *apoBa* mutants injected with *apoBb.1* MO (D,E) and *apoBb.1* mutants injected with *apoBa* MO (F) show dark yolk at 2 dpf (D) and complete absence of lipids in circulation at 3 dpf (E,F). **(G)** Kaplan–Meier curves showing significant decrease in *apoBb.1* mutant viability at 60 dpf (N=3, n_WT_=127, n_*apoBb.1*−/−_=130). **(H)** Decreased weight of *apoBb.1* homozygous mutants at 60 dpf, as compared to WT siblings (N=3, n_WT_ =87, n_*apoBb.1−/−*_=37). Scale bar: (A-F) 100μm, P Value: *<0.05.

**Supplemental Figure 3**. **(A)** Liver area in wt, *apoba*^−/−^*, apoBb.1*^−/−^ and *apoba*^−/−^*;apoBb.1*^*MO*^ embryos. **(B)** WISH at 24 hpf shows no differences in the expression of *foxa2* in *apoBb.1*^*MO*^ and *apoba*^−/−^*;apoBb.1*^*MO*^ embryos as compared to wt and *apoba*^−/−^ siblings. **(C)** H&E stained sections of 4 dpf embryos showing lipid accumulation in *apoBb.1*^−/−^ and *apoba*^−/−^*;apoBb.1*^−/−^ embryos. Arrowheads point to lipid droplets within hepatocytes. Scale bar: (C) 50μm. P Value: ***<0.001

**Supplemental Figure 4**: **(A)** Schematic representation of the zebrafish embryonic vasculature, with blue square marking the cranial vasculature (shown in F,G), and yellow square depicting the trunk vasculature (I,J,L). **(B-E)** Confocal images of *apoBb.1*^−/−^ at 2 (B,C) and 3 (D,E) dpf, uninjected (B,D) or injected with *apoBb.1* MO showing no defects caused by MO injection. (F,G) Dorsal view of the cranial vessels at 4 dpf showing the presence of ectopic vascular loops in the PCeV of *Tg(fli1:eGFP)*;*apoBa*^−/−^;*apoBb.1*^*MO*^ (arrowheads); quantified in (H) (N=3, n_*apoBa*−/−_=15, n_*apoBa*−/−*;apoBb.1MO*_=19). **(I,J)** Confocal images of the trunk of 5 dpf *Tg(fli1:eGFP);apoBa*^−/−^ (I) and *Tg(fli1:EGFP);apoBa*^−/−^*;apoBb.1*^*MO*^ (J) exhibiting ectopic angiogenic sprouts (arrowheads) arising in the ISVs, quantified in (K) (N=3, n_*apoBa*−/−_=14, n_*apoBa*−/−*;apoBb.1MO*_=21). **(L)** Confocal images of 2 dpf *Tg(fli:eGFP)*^*y1*^ embryo injected intravascularly with DiI-LDL, showing proper distribution of DiI-LDL in circulation. **(M)** Schematic model describing the experimental system for production of truncated ApoB fragments.Scale bar: (A-J) 50μm, (L) 30μm, P Value: *<0.05.

