## Supplementary figures and images for "Novel zebrafish mutants reveal new roles for Apolipoprotein B during embryonic development and pathological conditions"

### Supplemental Figure 1

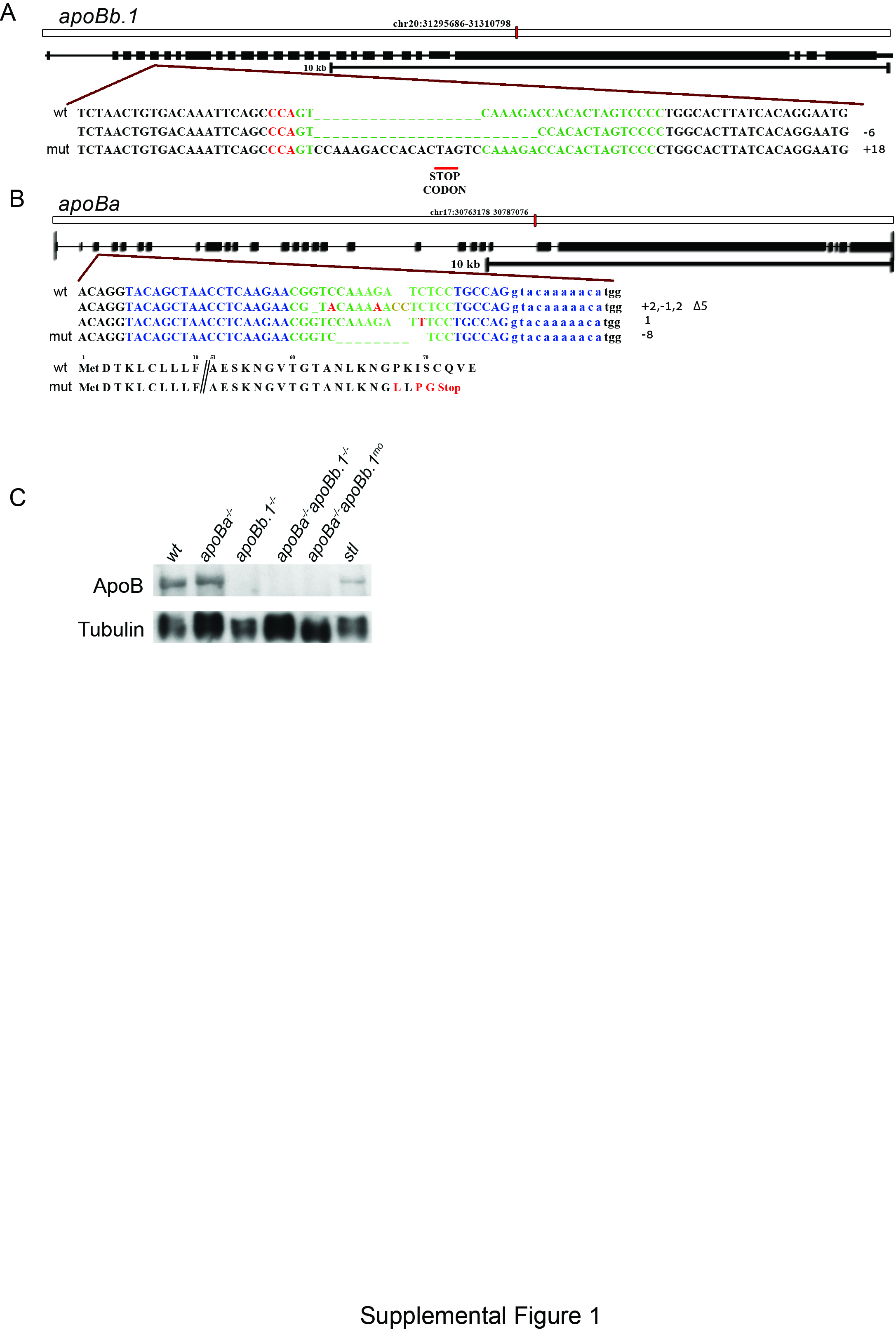

### Supplemental Figure 2

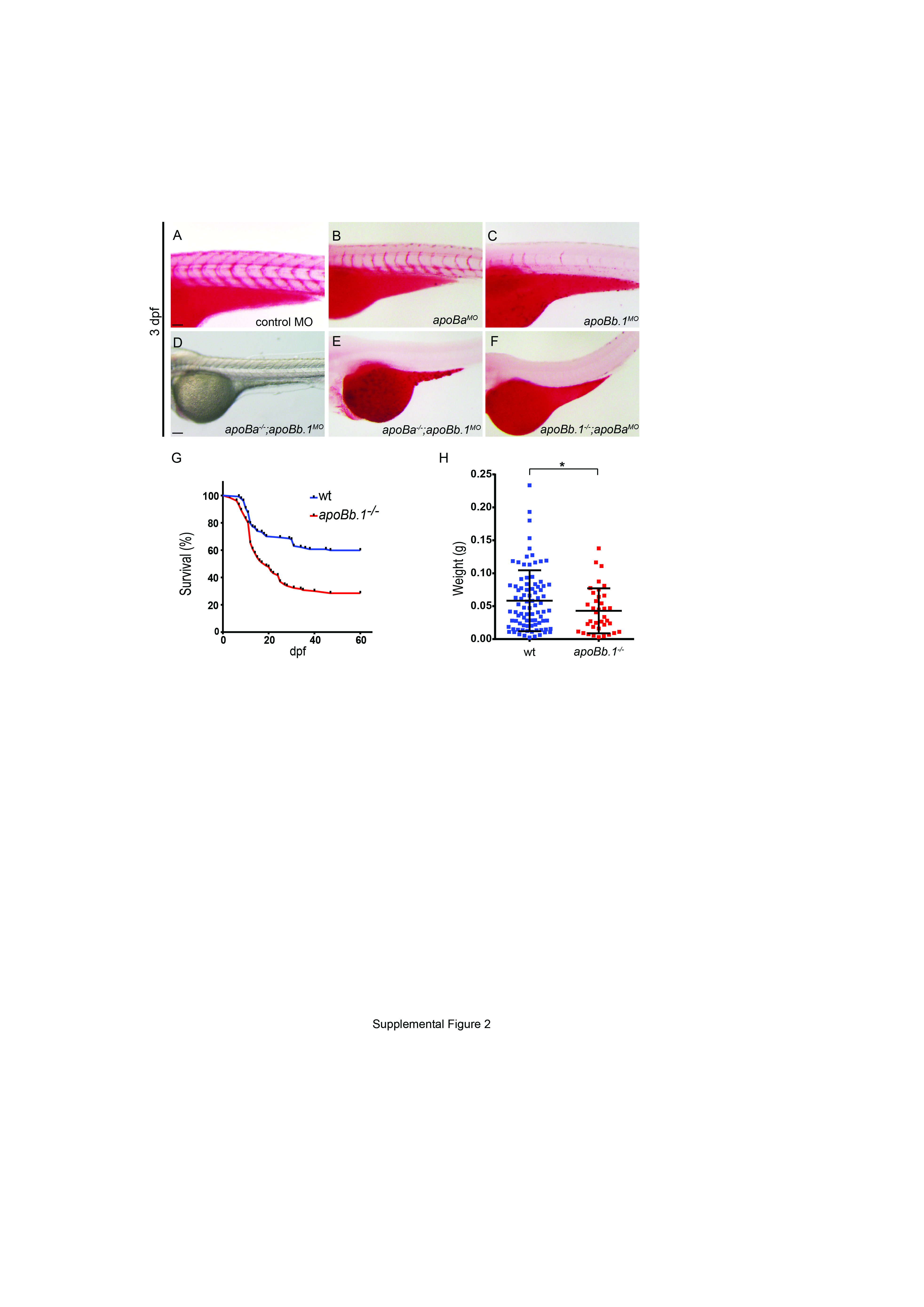

### Supplemental Figure 3

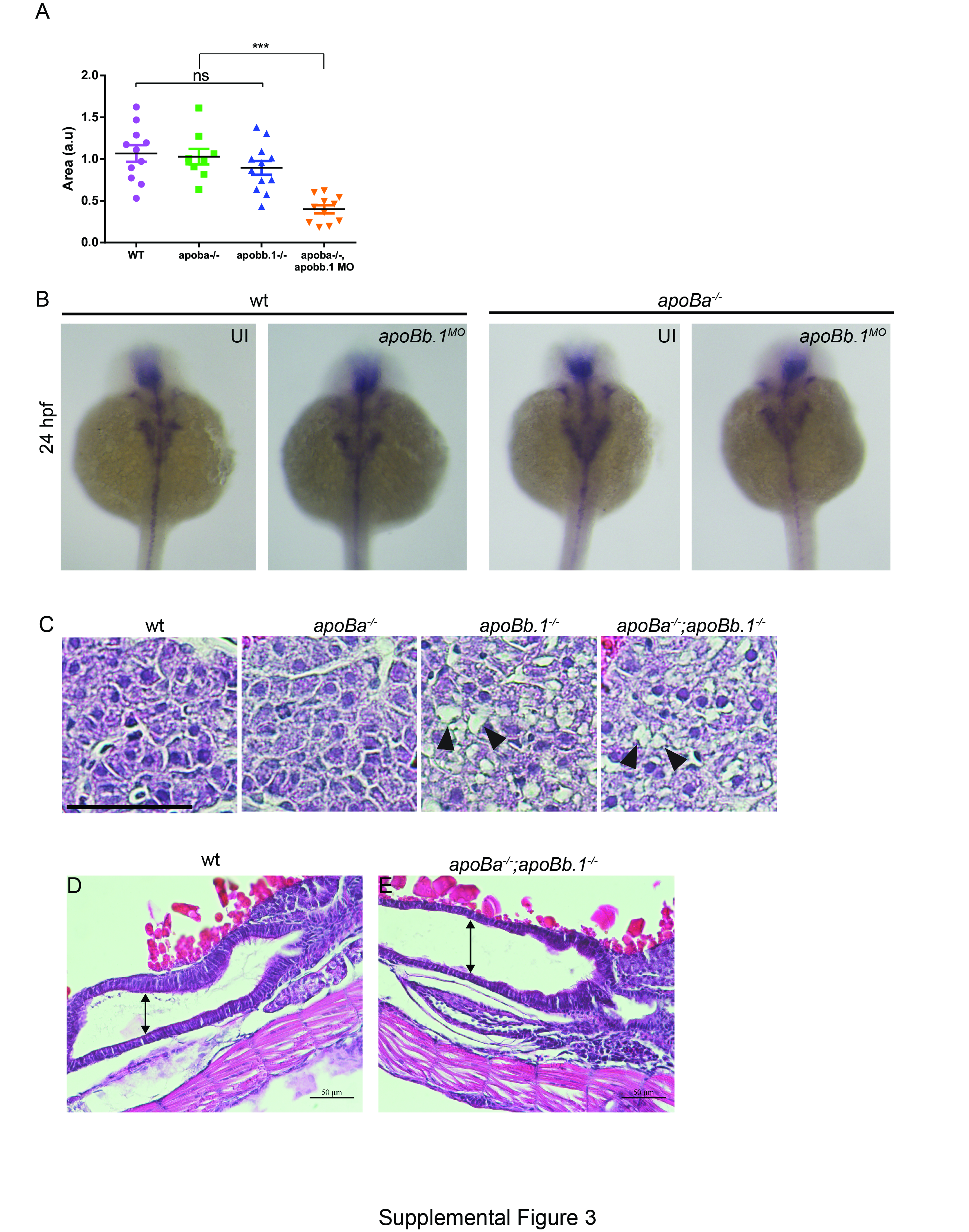

### Supplemental Figure 4

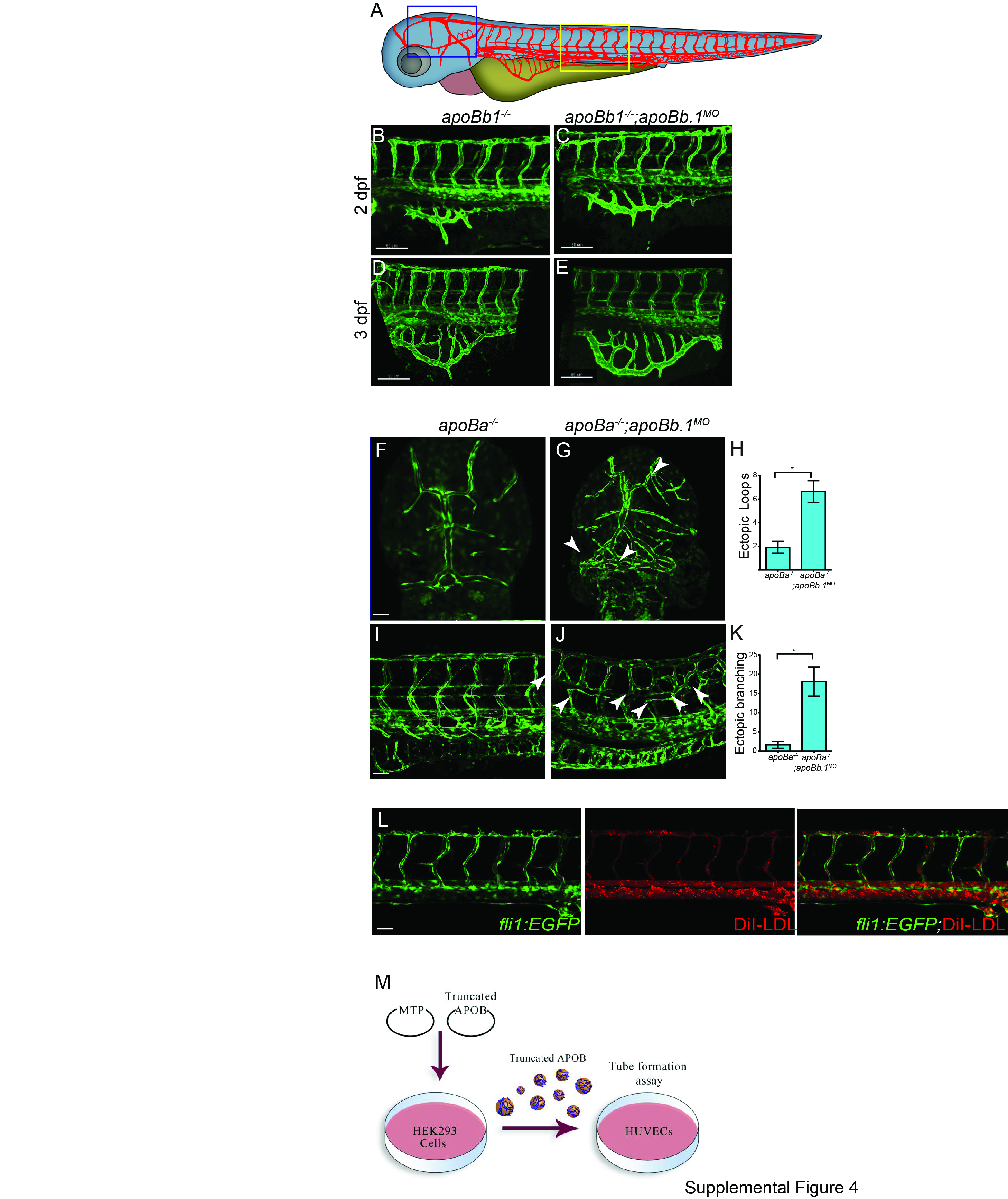
